# Activation of targetable inflammatory immune signaling is seen in Myelodysplastic Syndromes with SF3B1 mutations

**DOI:** 10.1101/2022.03.09.483595

**Authors:** Gaurav S Choudhary, Andrea Pellagatti, Bogos Agianian, Molly A Smith, Tushar D Bhagat, Shanisha Gordon-Mitchell, Sanjay Pandey, Nishi Shah, Srinivas Aluri, Leya Schwartz, Violetta Steeples, Robert N Booher, Murali Ramachandra, Maria Elena Samson, Milagros Carbajal, Kith Pradhan, Teresa V. Bowman, Manoj M Pillai, Britta Will, Amittha Wickrema, Aditi Shastri, Robert Bradley, Robert Martell, Ulrich G. Steidl, Evripidis Gavathiotis, Jacqueline Boultwood, Daniel T. Starczynowski, Amit Verma

## Abstract

**Background:** Mutations in the SF3B1 splicing factor are commonly seen in Myelodysplastic syndromes (MDS) and Acute Myeloid Leukemia (AML), yet the specific oncogenic pathways activated by missplicing have not been fully elucidated. Inflammatory immune pathways have been shown to play roles in pathogenesis of MDS, though the exact mechanisms of their activation in splicing mutant cases are not well understood.

**Methods:** RNA-seq data from SF3B1 mutant samples was analyzed and functional roles of IRAK4 isoforms were determined. Efficacy of IRAK4 inhibition was evaluated in pre-clinical models of MDS/AML

**Results:** RNA-seq splicing analysis of innate immune mediators in SF3B1 mutant MDS samples revealed retention of full-length exon 6 of interleukin-1 receptor-associated kinase 4 (IRAK4), a critical downstream mediator that links the Myddosome to inflammatory NF-kB activation. Exon 6 retention leads to a longer isoform, encoding a protein (IRAK4-Long) that contains the entire death domain and kinase domain, leading to maximal activation of NF-kB. Cells with wild-type SF3B1 contain smaller IRAK4 isoforms that are targeted for proteosomal degradation. Expression of IRAK4-Long in SF3B1 mutant cells induces TRAF6 activation leading to K63-linked ubiquitination of CDK2, associated with a block in hematopoietic differentiation. Inhibition of IRAK4 with CA-4948, leads to reduction in NF-kB activation, inflammatory cytokine production, enhanced myeloid differentiation in vitro and reduced leukemic growth in xenograft models.

**Conclusions:** SF3B1 mutation leads to expression of a therapeutically targetable, longer, oncogenic IRAK4 isoform in AML/MDS models.

## Introduction

Myeloid malignancies MDS and AML are associated with dismal prognosis and need newer therapeutic options. Mutations in spliceosome genes are commonly seen in MDS and AML and SF3B1 is the most frequently mutated gene in patients with MDS (1). Even though reports have shown that mutations in SF3B1 can lead to widespread changes in splicing (2), the exact pathways that are disrupted and lead to the pathogenesis of ineffective hematopoiesis and carcinogenesis in MDS and AML are not yet fully elucidated.

The innate immune signaling pathways have been shown to be activated in MDS and AML(3). The upstream activation of TLRs, IL1RAP and IL8 activate the IRAK/TRAF6 pathways and lead to NF-kB activation (4). Even though these pathways have been shown to be overactivated, a link between this activation and commonly seen SF3B1 mutation has not been established. We had recently shown that another splicing mutation in U2AF1 splicing factor can lead to retention of exon 4 leading to overexpression of IRAK4 in MDS and AML(5). In the present report, we now demonstrate that SF3B1 mutation leads to exon 6 retention leading to overexpression of active isoform of IRAK4 known as IRAK4-Long (IRAK4-L). We demonstrate that this IRAK4-L isoform containing the death domain can associate with MYD88 and lead to activation of downstream NF-kB signaling. These data demonstrate the link between SF3B1 mutation and oncogenic IRAK4 signaling in MDS/AML and also demonstrate the therapeutic potential of inhibiting this pathway via pharmacologic inhibitors.

## Results

### SF3B1 mutations are associated with inclusion of exon 6 in MDS

We sought to determine whether primary MDS samples with SF3B1 mutations have altered splicing of transcripts involved in immune and inflammatory pathways. Analysis of RNA-seq of purified CD34+ HSPCs from healthy controls and SF3B1 mutant MDS samples demonstrated an increased retention of full length exon 6 of IRAK4 in the SF3B1 mutant samples (Fig 1A). The full length retention of exon 6 was confirmed by RT-PCR in a larger cohort of primary samples demonstrating a significantly greater ratio of long isoform of IRAK4 in SF3B1 mutant samples when compared to healthy controls and MDS patients without splicing mutations (Fig 1B,C). To gain further insight into the regulation of *IRAK4* exon 6 inclusion in the presence of SF3B1 mutations, we generated an *IRAK4* exon 6 cassette splicing reporter in HEK293 cells (HEK293-*IRAK4*exon6). Transfection of mutant SF3B1-K700E into HEK293-*IRAK4*exon6 cells resulted in inclusion of *IRAK4* exon 6 from the splicing reporter compared to SF3B1-WT or vector-transfected HEK293-*IRAK4*exon6 cells (Fig 1D), indicating that SF3B1-K700E directly mediates inclusion of IRAK4 exon 6

**Figure 1.**
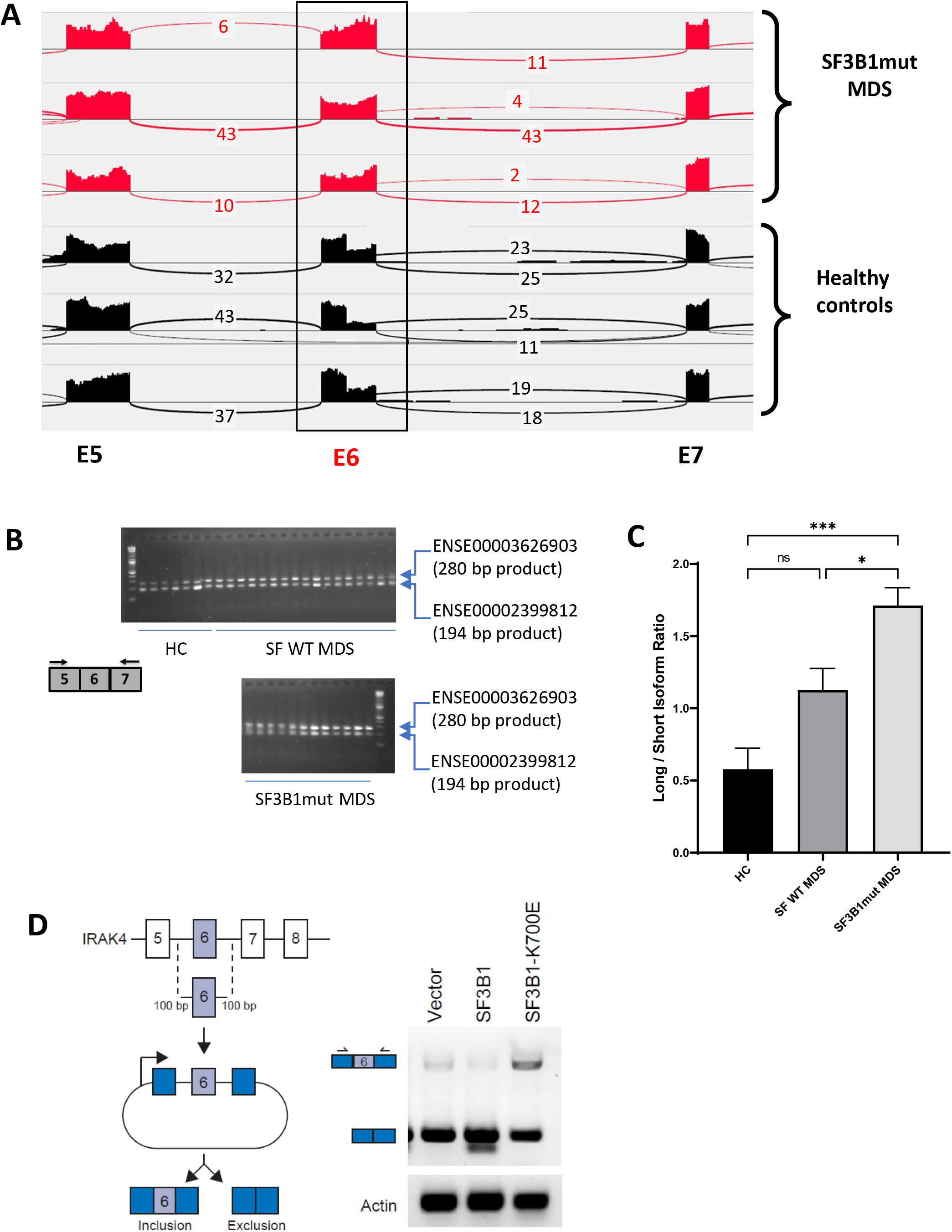
SF3B1 mutations are associated with inclusion of exon 6 in MDS and AML. A. Sashimi plots representing full length IRAK4 exon 6 inclusion or exclusion in CD34+ cells from healthy controls (n = 3), and from MDS patients with mutation in SF3B1 (MDS-SF3B1^mut^; n = 3) based on RNA-sequencing junction reads. B. RT-PCR analysis of CD34+ cells from healthy controls (HC, n = 7), from MDS patients with no splicing factor mutations (MDS-SF^WT^; n = 17), and from MDS patients with mutation in SF3B1 (MDS-SF3B1^mut^; n = 12) using primers flanking IRAK4 exon 6. C. Densitometric quantification of IRAK4 exon 6 inclusion calculated as the ratio of the long isoform versus the short isoform from panel B. Data represent the mean ± s.e.m. P-values were obtained using Kruskal–Wallis test with Dunn’s multiple comparisons test. * P<0.05, *** P<0.001 D. Schematic of IRAK4 exon 6 splicing reporter. IRAK4 exon 6 and 50 bp of flanking introns were cloned into a splicing reporter (pFlare5A)55 and stably expressed into HEK293 cells. SF3B1 and SF3B1-K700E were transfected and IRAK4 exon 6 splicing was measured by RT-PCR using primers within pFlare5A.

### SF3B1 mutation leads to longer, stable and functionally active isoform of IRAK4

To evaluate the effects of retention of full length exon 6 seen in SF3B1 mutant samples, we modelled the exon usage in vitro. Full length IRAK4 protein (IRAK4-L) is composed of a death domain, hinge region and a kinase domain (Fig 2A). Exon 6 encodes a part of the kinase domain of IRAK4 protein. The exclusion of mRNA sequence encoding amino acids 188-217 in exon 6 as seen in healthy controls can lead to two possible smaller IRAK4 isoforms. Isoform S1 (191 amino acids) could be produced due to formation of a premature stop codon (TGA) leading to a smaller IRAK4 isoform containing death, hinge and part of kinase domain. Isoform S2 (243 amino acids) (S2) could also be generated by utilizing an alternative translational start site (ATG) resulting in an IRAK4 isoform with only the kinase domain (Fig. 2A, Supplementary Fig. 1). We generated flag tagged plasmids of IRAK4 with full length exon 6 (IRAK4-L), and shorter S1 and S2 isoforms and transfected them in HEK-293T cells and evaluated their effects on NF-kB pathway activation. Immunoblotting analysis showed that plasmid with the full length exon 6 led to stable expression of IRAK4-long protein and resulted into increased phosphorylation of p65 when compared to the smaller isoforms (Fig 2B). By contrast, transfection of IRAK4 S1 and S2 constructs led to generation of lower molecular weight IRAK4 proteins with decreased expression when compared to the longer isoform (Fig 2B). Treatment with proteasome inhibitor (MG132) led to increased expression of the smaller IRAK4 bands demonstrating that these were targeted for proteasomal degradation (Fig 2C). Immunoprecipitation in HEK293T cells transfected with IRAK4-L, S1, S2 and HA-tagged Lys48 linked in presence of proteasome inhibitor MG132 demonstrated an increased association of Lys48 linked ubiquitin chains with shorter IRAK4 isoforms, validating ubiquitin mediated degradation. (Fig 2D).

**Figure 2:**
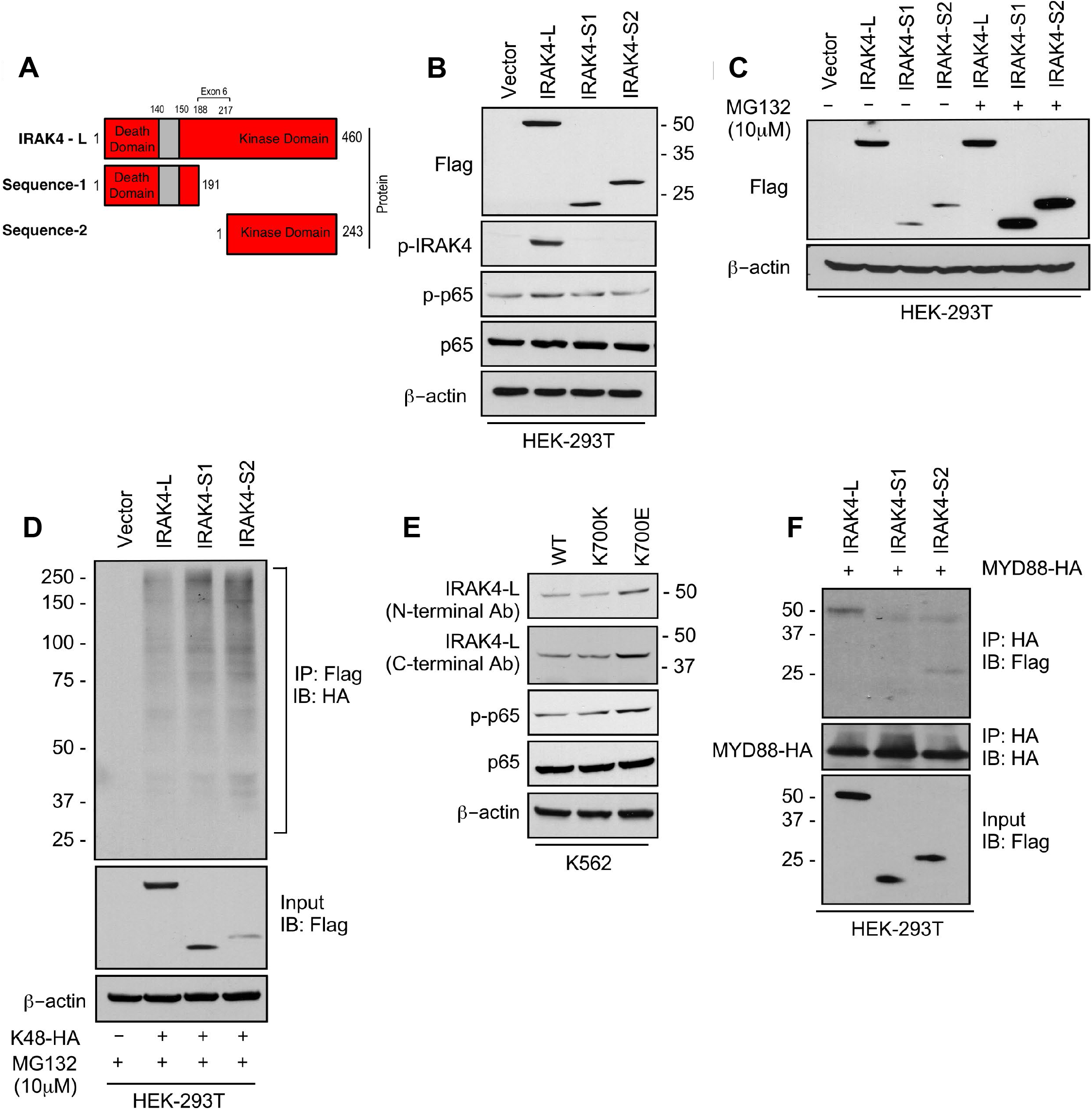
SF3B1 mutation leads to longer, stable and functionally active isoform of IRAK4. A. Schematic showing that IRAK4 protein consists of a kinase domain and death domain that associates with MYD88. The segment of exon 6 that is not included in WT controls encodes amino acids 188-217 and can lead to two smaller isoforms by generation of premature stop codon or use of alternative translational start site. B. HEK-293T cells were transfected with Flag tagged plasmids of IRAK4-long isoform containing full length exon 6, IRAK4-S1 and IRAK4-S2. p-IRAK4, p-p-65 and p65 were determined by immunoblotting. C. Flag tagged plasmids of IRAK4-long isoform containing full length exon 6, IRAK4-S1 and IRAK4-S2 were transfected in HEK293T cells and treated with proteasome inhibitor MG132 (10uM) overnight followed by immunoblotting. IRAK4-S1 and IRAK4-S2 constructs that led to smaller IRAK4 protein bands that were underexpressed and increased upon proteasomal inhibition. D. Ubiquitination of ectopic IRAK4 was determined in HEK-293T cells transfected with indicated plasmids in presence of MG132 (10uM) by immunoprecipitating with HA-specific antibody and immunoblotting with Flag. E. Immunoblotting analysis for indicated proteins in isogenic K562 cells with WT and K700E mutation of SF3B1. F. HEK-293T cells were transfected with Flag tagged IRAK4 isoforms and HA tagged MYD88. MYD88 was immunoprecipitated with HA-specific antibody and its association with IRAK4 was probed by immunoblotting with specific antibodies.

Next, we wanted to determine whether expression of the SF3B1 mutation leads to changes in isoforms and downstream NF-kB activation. Isogenic cells with WT and K700E mutation of SF3B1 were evaluated and the mutant cells demonstrated overexpression of IRAK4-L and higher activation of p-p65 (Fig 2E). Finally, we showed that the IRAK4-long containing the death domain was able to associate with MYD88 in immunoprecipitation experiments, while the shorter isoforms did not; thus providing the mechanistic basis of activation seen only with the IRAK4-long isoform (Fig 2F).

### CDK2 is ubiquinated at K63 residues downstream of IRAK4/TRAF6 in SF3B1 mutants

Having demonstrated overexpression of the active IRAK4-long isoform in SF3B1 mutant samples, we next wanted to determine the downstream pathways that are activated by these changes. Signal transduction via toll-like receptors is primarily mediated by IRAK4-TRAF6 activation and TRAF6 is an E3 ligase that conjugates ubiquitin chains to itself and its substrates. The consensus sequence of TRAF6 binding motif is P(P-2)XE(P0)XX(Acidic/Aromatic)(P+3) and is present in known TRAF6 substrates (Fig. 3A). Since MDS/AML are characterized by ineffective hematopoiesis due to block in myeloid differentiation, we performed a search for the TRAF6 motif in cell cycle regulators and determined that CDK2 could be a potential substrate (Fig 3A). Structural modeling of TRAF6 and CDK2 peptide show that positions P-2, P0 and P3 in CDK2 correspond to three distinct sub-pockets in the binding site of TRAF6 and supports the notion that CDK2 can be ubiquitinated by TRAF6 (Fig. 3B and Supplementary Figure 2). To directly test the role of TRAF6 dependent ubiquitination of CDK2 in chronic innate immune signaling, we ectopically expressed CDK2, IRAK4, and TRAF6 in presence of either Lys48 linked ubiquitin or Lys63 linked ubiquitin (Ub).CDK2 was associated with a much stronger K63 Ub that was dependent on TRAF6 and IRAK4 overexpression (Fig 3C). To validate this result, we generated alanine and glycine mutants (CDK2 Myc 122 and CDK2 Myc 123) within CDK2 binding site of TRAF6 and examined their ubiquitination in presence of IRAK4-L and TRAF6 by immunoprecipitating CDK2. Immunoblotting analysis show that TRAF6 mediated Lys63 linked ubiquitination was decreased in CDK2 mutants as compared to wild type CDK2 (Fig. 3D).

**Figure 3:**
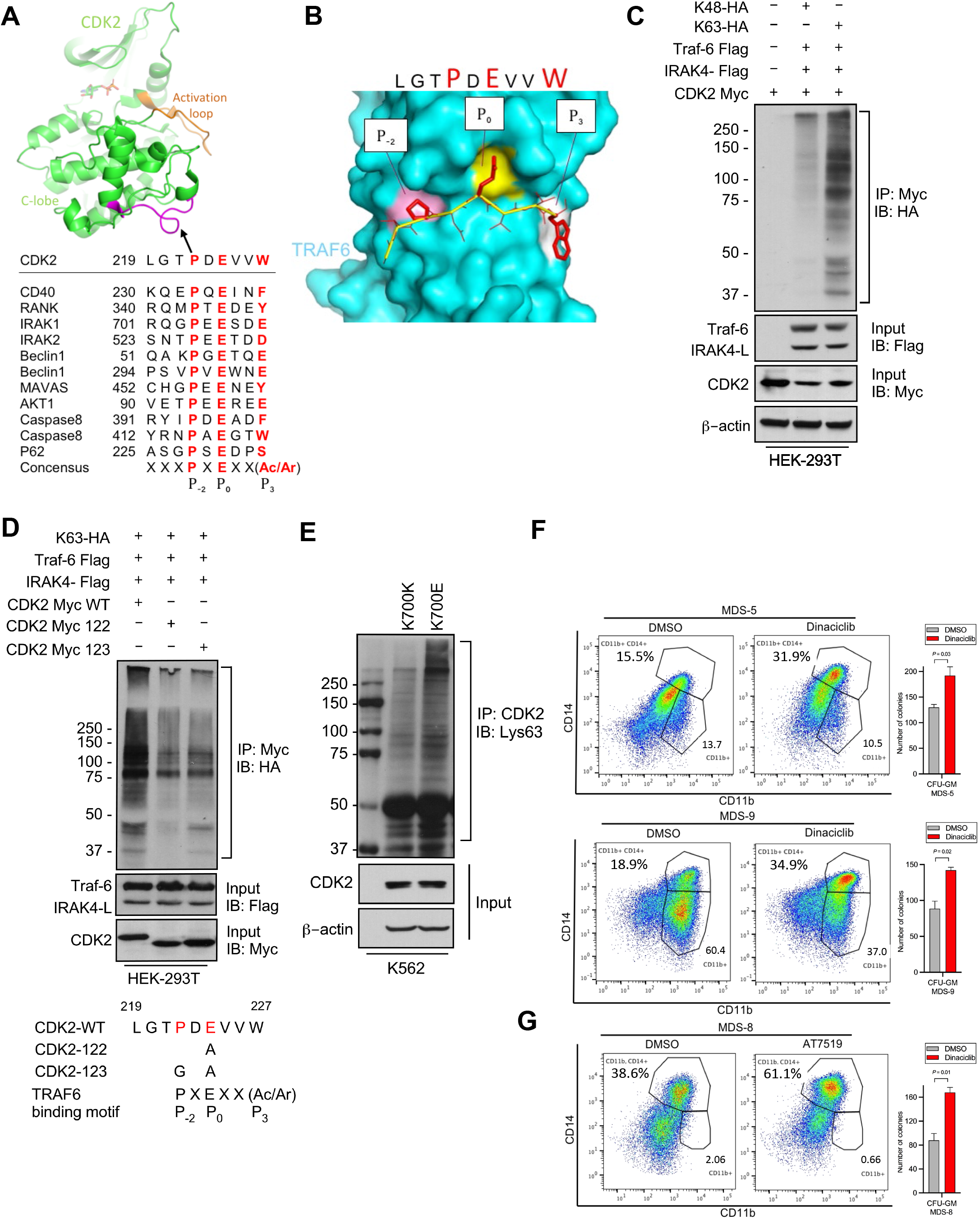
IRAK4/TRAF6 signaling regulates CDK2 ubiquitination in SF3B1 mutant cells. A. CDK2 structure (PDB: 1GY3) bound to ATP is shown in cartoon representation. The putative TRAF6 binding peptide (magenta) is located at a loop between αG and αH in the C-lobe of the kinase. Sequence alignment of CDK2 peptide with common binding sequences with TRAF6. The consensus pattern with conserved P_-2_, P_0_ and P_3_ positions is shown at the bottom. B. MM-GBSA docking of CDK2 peptide to substrate binding domain of TRAF6 (PDB:1LB5). The top 1 populated cluster is shown. TRAF6 and CDK2 peptide are shown in ribbon and surface representation, respectively. Positions P-2, P0 and P3 correspond to three distinct sub-pockets in the binding site, formed largely by residues Y473, M450, G472 (pink), A458, G470, K469 (yellow) and H376, V374, R392 (white). Capital lettering of conserved residues P, E, W within the shown peptides depicts successful docking in corresponding sub-pockets. C. Lys63-linked proteins were immunoprecipitated in K562 cells with WT and K700E SF3B1 mutation and CDK2 ubiquitination was determined by CDK2 specific antibody. D. Ubiquitination of CDK2 was determined in HEK-293T cells transfected with indicated plasmids by immunoprecipitating with HA specific antibody and immunoblotting with either Lys48 or Lys63 antibody. E. Lys63-linked proteins were immunoprecipitated in HEK-293T cells transfected with indicated plasmids and CDK2 ubiquitination was determined by Myc-tag specific antibody. Sequence alignment of optimum amino-acids sequence predicted for TRAF6 with wild type (WT) and mutant CDK2 (CDK2-122 and CDK2-123) F,G:MDS patient derived samples (Bone marrow/ Peripheral Blood) were treated with DMSO or Dinaciclib (10nM) or AT7519 (10nM) for 14 days on methylcellulose clonogenic assays. The samples were evaluated for colony formation CFU-GM (colony forming unit-granulocyte macrophage) and for myeloid differentiation on colonies that were subjected to flow cytometry.

Furthermore, we used isogenic cell line containing K700E mutant and K700K control SF3B1 and demonstrated that the mutant cells contained heavily K63 Ub CDK2 (Fig 3E). Since K63 ubiquitination leads to increased protein stability and interactions, we tested whether inhibiting CDK2 would lead to functional effects in patient samples containing the SF3B1 mutation. Cases of MDS with SF3B1 mutations are characterized by ineffective hematopoiesis and treatment with CDK2 inhibitors in vitro led to increased hematopoietic colony formation and increased myeloid differentiation as evident by CD14 expression seen in 3 primary samples. (Fig 3F).

### IRAK4 inhibition leads to increased differentiation in SF3B1 mutant MDS

The hallmark of MDS and AML is a block in differentiation leading to dysplastic maturation and cytopenias. We wanted to evaluate the efficacy of IRAK4 inhibition in SF3B1 mutated MDS with a clinically relevant inhibitor, CA4948, which binds to the kinase domain on the protein (Fig 4A). CA4948 treatment is able to inhibit downstream NF-kB activation upon TLR engagement in leukemic THP1 cells (Fig 4B). This IRAK4 inhibitor also led to reduction in cytokine production in leukemic cells after engagements of Myd88/TLR pathways (Fig 4C). Bone marrow mononuclear cells from MDS patients with SF3B1 mutation were cultured in the presence and absence of the IRAK4 inhibitor in vitro and then colonies were assessed by flow cytometry for myeloid differentiation. We observed a significant increase in myeloid colonies as well as increase in myeloid differentiation markers after IRAK4 inhibition in 3 distinct samples (Fig 4D). Treatment of healthy human CD34 stem and progenitor cells with IRAK4 inhibitor did not lead to any significant changes in colony numbers (Fig 4E). We also performed IRAK4 knockdown with siRNAs in a primary MDS sample with SF3B1 mutations and observed an increase in colonies and myeloid differentiation upon knockdown, validating IRAK4 as a target for relieving the differentiation block seen in MDS (Fig 4F).

**Figure 4:**
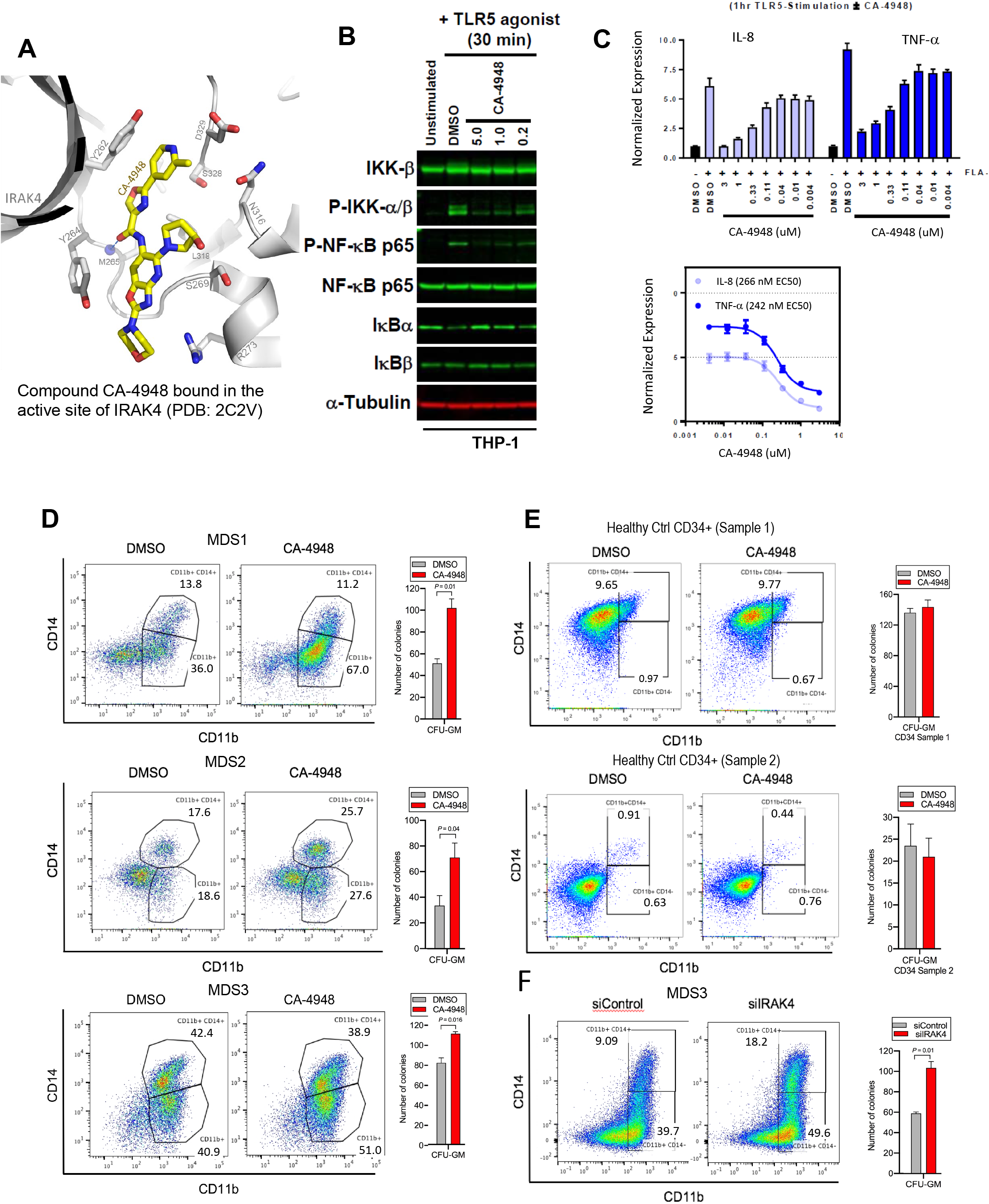
IRAK4 inhibition with CA-4948 promotes differentiation in MDS/AML. A. Crystal structure of CA-4948 bound in the to active site of IRAK4 (PDB: 7C2V). Interacting residues of IRAK4 are shown in sticks. B. Immunoblotting analysis for indicated proteins in THP-1 cells that were stimulated by TLR-5 and treated either with DMSO or CA-4948. C. mRNA levels of IL-8 and TNF-α in OCI-AML2 cells stimulated by TLR5 ligand and were either treated with DMSO or CA-4948. D. MDS patient derived samples with SF3B1 mutation were treated with IRAK4 inhibitor (CA4948) or control in methylcellulose clonogenic assays and the analyzed for myeloid colony formation. Colonies were picked and analyzed by FACS for myeloid differentiation markers CD11b and CD14. E. Healthy CD34+ stem and progenitor cells were grown in clonogenic assays with and without IRAK4 inhibitor (CA4948). Myeloid colonies were counted at Day 14 and colonies were analyzed by FACS F. MDS patient sample with SF3B1 mutation was treated with siRNAs against IRAK4 and control and grown in clonogenic assays. The sample were evaluated for myeloid colony formation and for differentiation by analyzing their CD11b and CD14 expression with flow cytometry.

### IRAK4 overexpression in MDS is associated with worse prognostic features and its inhibition leads to reduction in MDS/AML clones in vivo

Next we tested whether overexpression of IRAK4-long was associated with adverse clinical features in a large cohort of MDS samples (N=183) (6). IRAK4-long expression was determined from an existing expression dataset from selected CD34+ cells and used to divide the cohort into high and low IRAK4 based on median expression. We observed that cohort with higher IRAK4-long expression had lower platelet counts, higher RBC transfusion needs and higher leukemic blast counts, all indicative of worse clinical features (Fig 5A). Next, we established xenografts with MDS samples with SF3B1 mutation and treated the NSG mice with either IRAK4 inhibitor (CA4948) and placebo controls. Treatment with IRAK4 inhibitor led to decrease in MDS cells after 3-4 weeks of treatment in 5 distinct samples (Fig 5B-D, Supplementary Fig 3).

**Figure 5:**
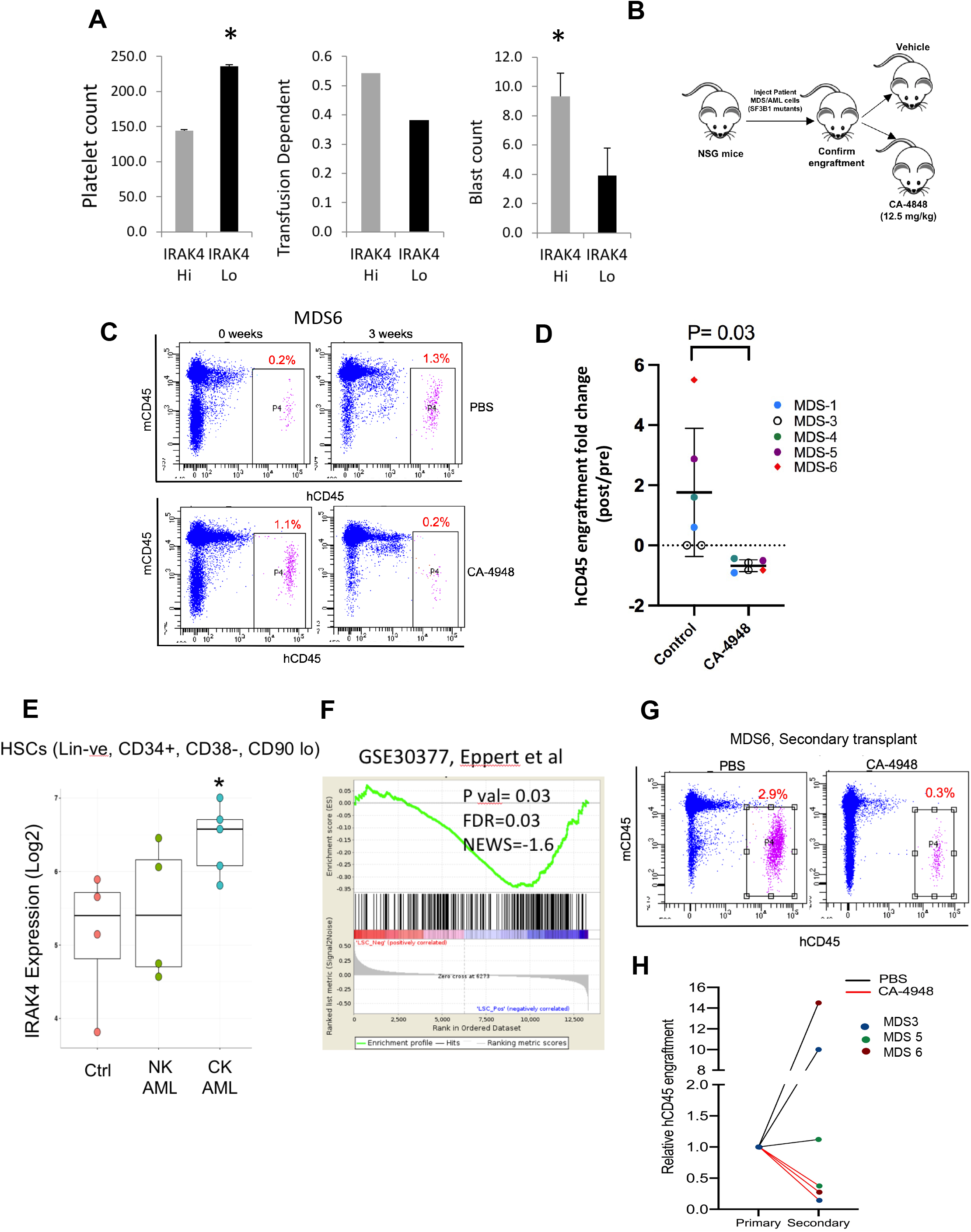
IRAK4 overexpression is associated with adverse clinical features in human MDS and its inhibition leads to reduction in MDS clones in vivo. A. IRAK4 expression in 183 samples from MDS patient bone marrow CD34+ cells was correlated with platelet counts, RBC transfusion dependency and Blast counts. Cohort with high IRAK4 expression (> Median) were associated with lower platelet counts, higher transfusion dependence and higher blast counts (Ttest, P Value< 0.05). B. NSG mice were xenografted with MDS cells with SF3B1 mutation. After engraftment, mice were treated with either vehicle or IRAK4 inhibitor (CA-4849, 12.5 mg/kg) and bone marrow aspirates were used to evaluate human cells engraftment by flow cytometry. C,D. Summary of human cell engraftment for xenografted mice treated with either IRAK4 inhibitor (CA-4948) or vehicle. Representative MDS sample xenograft with SF3B1 K700E mutation is shown E. IRAK4 expression in sorted HSCs (Lineage –ve, CD34+, CD38-) cells from MDS/AML cases (N=9) and Controls (N=5). Significantly increased IRAK4 expression is seen in samples with complex karyotype (CK) when compared to healthy controls (TTest, P<0.05). F. Gene expression signature associated with high IRAK4 expression (>median, from set of 183 MDS CD34+ cells) was compared to know leukemic stem cell signature and showed significant similarity by GSEA analysis. G,H. MDS bone marrow cells were isolated and purified from mice either treated with vehicle or CA-4948 in and then xenografted in secondary recipient NSG mice. Human cell engraftment was evaluated after 4 weeks by flow cytometry on bone marrow aspirates.

Since innate immune signaling pathways have been shown to play roles in leukemic stem cell propagation (3), we next evaluated IRAK4-long expression in a dataset derived from sorted AML/MDS HSCs and healthy controls (7). IRAK4-Long was found to be elevated in selected cases of AML HSCs, most significantly in adverse risk patients with complex cytogenetics (Fig 5E). Consistent with this finding, we derived a transcriptomic signature of IRAK4 overexpression from 183 MDS samples (Supplementary Fig 4) and compared it to a published leukemic stem cell signature (8) and found significant concordance (Fig 5F). Lastly, we tested the efficacy of IRAK4 inhibition in disease initiation in vivo. IRAK4 inhibitor and placebo treated xenograft mice were sacrificed and cells used for secondary xenografts. IRAK4 inhibition led to reduced MDS clones in secondary transplants, demonstrating decrease in disease initiating activity (Fig 5G,H, Supplementary Fig 5, Summarized in Supplemental Fig 6).

## Discussion

Myelodysplastic Syndromes (MDS) and acute myeloid leukemia (AML) are frequently incurable hematologic disorders characterized by cytopenias that are a major cause of morbidity and mortality. Mutations in splicing genes are one of the most common alterations in MDS and AML and predominantly affect SF3B1, U2AF1 and SRSF2 (1). SF3B1 is the most commonly mutated splicing gene in MDS and can also be mutated in solid tumors such as breast cancer, melanoma and others.(9) Even though mutations in SF3B1 can affect splicing of numerous genes, the exact pathways that are disrupted by aberrant splicing and lead to ineffective hematopoiesis are not well elucidated. We recently demonstrated that mutant U2AF1 (S34F) directly regulates IRAK4 exon 4 retention in MDS/AML, which results in expression of the longer IRAK4 isoform (IRAK4-L)*(5)* In the present report, we demonstrate that SF3B1 mutations also lead to production of active IRAK4-L isoforms but by aberrant retention of exon 6 (Supplemental Fig 6). The functional activation of the IRAK4 pathway by two different exon retention events by U2AF1 mutations and SF3B1 mutations suggest that this is a very important functional event in MDS pathobiology.

While IRAK4 and its downstream pathways have been studied in innate immunity and in the context of inflammatory diseases, its molecular and cellular effects in MDS/AML are relatively understudied. Recent work has shown that innate immune signaling mediators such as Interleukin receptor 1 receptor accessory protein (IL1RAP), Interleukin8/CXCR2, Toll like receptors (TLRs) as well as myeloid derived suppressor cells are activated in MDS (7,10–13). Our findings further support the role of activated innate immune signaling pathways in MDS and suggest that interaction of IRAK4 with upstream TLR/MyD88 pathways is critical in MDS pathogenesis. Demonstration of overexpression of active IRAK4 isoforms in splicing mutant MDS suggests that these cells are primed to respond to upstream TLR and MyD88 activation. In fact, since active IRAK4 isoforms are expressed in different spliceosome mutant subsets, they potentially represents a shared downstream functional pathway that regulates MDS/AML malignant cell survival.

CA-4948 is a specific inhibitor of IRAK4 that is now being tested in clinical trials in MDS/AML and Lymphomas (ClinicalTrials.gov: NCT04278768; NCT03328078) and in preliminary findings has shown a tolerable safety profile. Early clinical data appears encouraging in spliceosome mutant patients, supporting the validity of IRAK4 as a therapeutic target in SF3B1 mutant MDS/AML (14). Our data suggests that inhibiting IRAK4 pharmacologically leads to shrinkage of the MDS/AML clones in vivo and this correlates with decreased blast counts seen with CA4948 monotherapy in relapsed/refractory MDS/AML in early results. MDS/AML are characterized by block in hematopoietic differentiation that leads to neutropenia and subsequent infections. Our data suggests that IRAK4 inhibition promotes myeloid differentiation in SF3B1 mutant cases. This affect can be therapeutically advantageous in clinical trials and supports further testing.

## Methods

(Study Schema outlined in Supplemental Fig 7)

### Cell lines and human samples

HEK-293T and THP1 were purchased from the American Type Culture Collection. MDS-L was provided by Dr. Kaoru Tohyama (Kawasaki Medical School, Okayama, Japan)(15,16). Cell-lines were cultured according to manufacturer’s instruction in either RPMI 1640, IMDM or DMEM medium supplemented with 10% FBS. MDS-L cell line was cultured in the presence of 10 ng/mL human recombinant IL-3 (Stem Cell Technologies). The MDS/AML patient samples used in this study were obtained with written informed consent under approval by the IRBs of the Albert Einstein College of Medicine. K562 isogenic cells with WT and K700E mutation of SF3B1 were purchased from Horizon Discovery.

### Plasmids and Reagents

The IRAK4 Inhibitor CA-4948 was received from Curis Inc. Dinaciclib and AT7519 was purchased from Selleck Chemicals. Recombinant human IL-1β and IL-3 was purchased from PeproTech. TLR-5 was purchased from Invivogen. IRAK4-S1 and IRAK4-S2 constructs were obtained from Integrated DNA Technologies IDT and cloned into pCDNA3.1 plasmid. IRAK4-L-Flag, pCNDA3.1, MYD-88-HA, Traf6-Flag was provided by Dr. Daniel Starczynowski (Cincinnati Children’s Hospital Medical Center, Cincinnati). K48-HA, K63-HA, CDK2-Myc was purchased from Addgene. The plasmids were transfected using Lipofectamine™ 2000 Transfection Reagent (Thermo Fisher Scientific) according to manufacture’s instructions. CDK2 mutants were generated using QuikChange II XL Site-Directed Mutagenesis Kit (Agilent Technologies) siControl and siIRAK4 were purchased from Horizon Discovery and transfected into human AML samples using Amaxa Nucleofector Kit T (Lonza) (program number G-016) according to manufacturer’s protocol.

### RNA sequencing of MDS CD34+ cells

Total RNA from MDS and control bone marrow CD34+cells was extracted using TRIzol with a linear acrylamide carrier, treated with DNase I (Life technologies) and purified using Agencourt RNAClean XP beads (Beckman Coulter). RNA quality was determined using an RNA 6000 Bioanalyzer pico kit (Agilent). A cDNA library was produced using a SMARTer library preparation protocol (Clontech). Sequencing (100-bp paired-end reads) was performed on an Illumina HiSeq4000. The reads were mapped to human genome GRCh37 using HISAT2 version 2.0.0-beta. Uniquely mapped read pairs were counted using htseq-count. The data have been deposited in the NCBI’s Gene Expression Omnibus (GEO) repository (accession number: GSE114922).

### Immunoblotting and Immunoprecipitation

Total protein lysates were obtained from cells by lysing the samples in 1% NP-40 lysis buffer 20 mmol/l Tris-HCl, pH 7.5; 1 mmol/l EDTA; 150 mmol/l NaCl; 1% NP-40), containing phosphatase inhibitor cocktails 2 and 3 and protease inhibitors (Sigma-Aldrich), for 30–45min at 4 °C. Equal amount of protein was prepared by calculating protein concentration using Bradford reagent (Bio-Rad) and 50μg of protein was resolved on 10–12% SDS-PAGE followed by transferring to nitrocellulose or PVDF membranes (EMD-Millipore). For immunoprecipitation, cells were lysed in 2% CHAPS lysis buffer (20 mmol/l Tris-HCl, pH 7.5; 150 mmol/l NaCl; 1 mmol/l EDTA; 2% CHAPS; Sigma-Aldrich) with protease and phosphatase inhibitors (Sigma-Aldrich) for 1 hour on ice. Protein lysates were incubated with primary antibody overnight at 4 °C. Protein A agarose beads ((Rockland Immunochemicals, Gilbertsville, PA) were added equally to all the samples and incubated for 1hr 4 °C. The beads were washed three times with CHAPS, eluted with loading buffer supplemented with 2-mercaptoethanol (Sigma-Aldrich) and western blotting was performed. In ubiquitination experiments, MG132 and N-ethylmaleimide (Sigma-Aldrich) were added to lysis buffer with dithiothreitol (Sigma-Aldrich) to prepare lysates. Western blot analysis was performed with the following antibodies: IRAK4 C-term (ab5985; Abcam), IRAK N-term (4363; Cell Signaling Technology), Flag (F3165; Sigma), IRAK1 (sc-7883; Santa Cruz), phospho-IRAK1 (T209) (A1074; AssaybioTech), IKKβ (2370; Cell Signaling Technology), phospho-IKKα/β (2697; Cell Signaling Technology), p38 MAPK (9212; Cell Signaling Technology), phopho-p38 MAPK (4631; Cell Signaling Technology), ERK (4695; Cell Signaling Technology), phospho-ERK (4377; Cell Signaling Technology), p-JNK (4668; Cell Signaling), p65 (8242; Cell Signaling Technology), phospho-p65 (3033; Cell Signaling Technology) and b-actin (Santa-Cruz Biotechnology).

### Flow Cytometry

The antibodies used for flow cytometric analysis of human MDS and AML cells included Mouse CD45 FITC; Human CD45 PeCy7 (e-Bioscience); Human CD4-Pacific Orange; Human CD8-Pacific Orange; Human CD19-Pacific Blue; Human CD20-Pacific Blue; Human CD11b-APC (Thermo Fisher); Human CD34-PE (Miltenyi Biotec);and Human CD33-APC-Cy7 (Abcam).

### Clonogenic progenitor assays

For clonogenic assays with primary MDS/AML cells, primary patient samples and healthy controls were plated in methylcellulose (Stem Cell Technologies H4435, Vancouver, CA) with the CA-4948 and control, and colonies were counted after 14-17 days.

### Xenografts

This study was performed in strict accordance with the recommendations in the Guide for the Care and Use of Laboratory Animals of the National Institutes of Health. All of the animals were handled according to approved institutional animal care and use committee (IACUC) protocols (001371) of the Albert Einstein College of Medicine. Mice were sacrificed and assessed for tumor burden measurements. For primary patient-derived xenografts NOD/SCID IL2Rgamma KO NSG mice were irradiated (200 rads) 24 hours prior to injection. Mononuclear cells from primary MDS patients were isolated by Ficoll separation. 2-5 x 10^6^ MNCs were administered via tail vein injection. 3-4 weeks later, BM aspiration were performed and analyzed by flow cytometry for the human cell engraftment. Mice were considered to be engrafted if they showed 0.1% or higher human derived CD45+ cells. The engrafted mice were randomized for treatment with 12.5 mg/kg/d of CA-4948 or control 5 times a week for indicated times. Following treatment, BM aspirations and flow cytometry analysis for the above-mentioned markers were performed. All mice were bred, housed and handled in the animal facility of Albert Einstein College of Medicine.

### Reporter cell lines

IRAK4 exon 6 and its flanking intronic sequences were inserted into the EcoRI and BamHI sites of pFlareA vector. pFlareA-exon4 vectors were linearized by DraIII and transfected into 293T cells. Cell colonies stably expressing the reporter were selected by 1 mg/mL G418 and stable expression of GFP and RFP.

### PCR

Primer3 designed primers

Fw - TGAACGACCCATTTCTGTTGG

Rev - GAGTCTGTCTAGCAATGAACCA

Expected product sizes:

Exon 6 short isoform – 194 bp

Exon 6 long isoform – 280 bp

cDNA was generated using a High-Capacity cDNA Reverse Transcription Kit (Thermo-Fisher). The primers listed above were used with Maxima Hotstart PCR mastermix to amplify the region containing exon 6 of IRAK4 and PCR products separated by electrophoresis on a 2 % agarose gel.

### Peptide Docking

3D structure of CDK2 peptide was created using BioLuminate (Schrodinger LLC, New York). The peptide was docked to TRAF6 substate binding domain (PDB code: 1LB5) using Glide (Schrodinger LLC) using “SP Peptide” mode that ensures enhanced conformational sampling of flexible polypeptides (17). To assure more accurate pose ranking, docked peptide poses were scored using implicit solvent MM-GBSA calculations. Final MM-GBSA interaction energies were used to create and rank peptide pose clusters.

## Data Availability

Publicly available dataset was used (GSE114922) for splicing isoforms in MDS.

## Supplementary Data

**Supplementary Figure 1:**
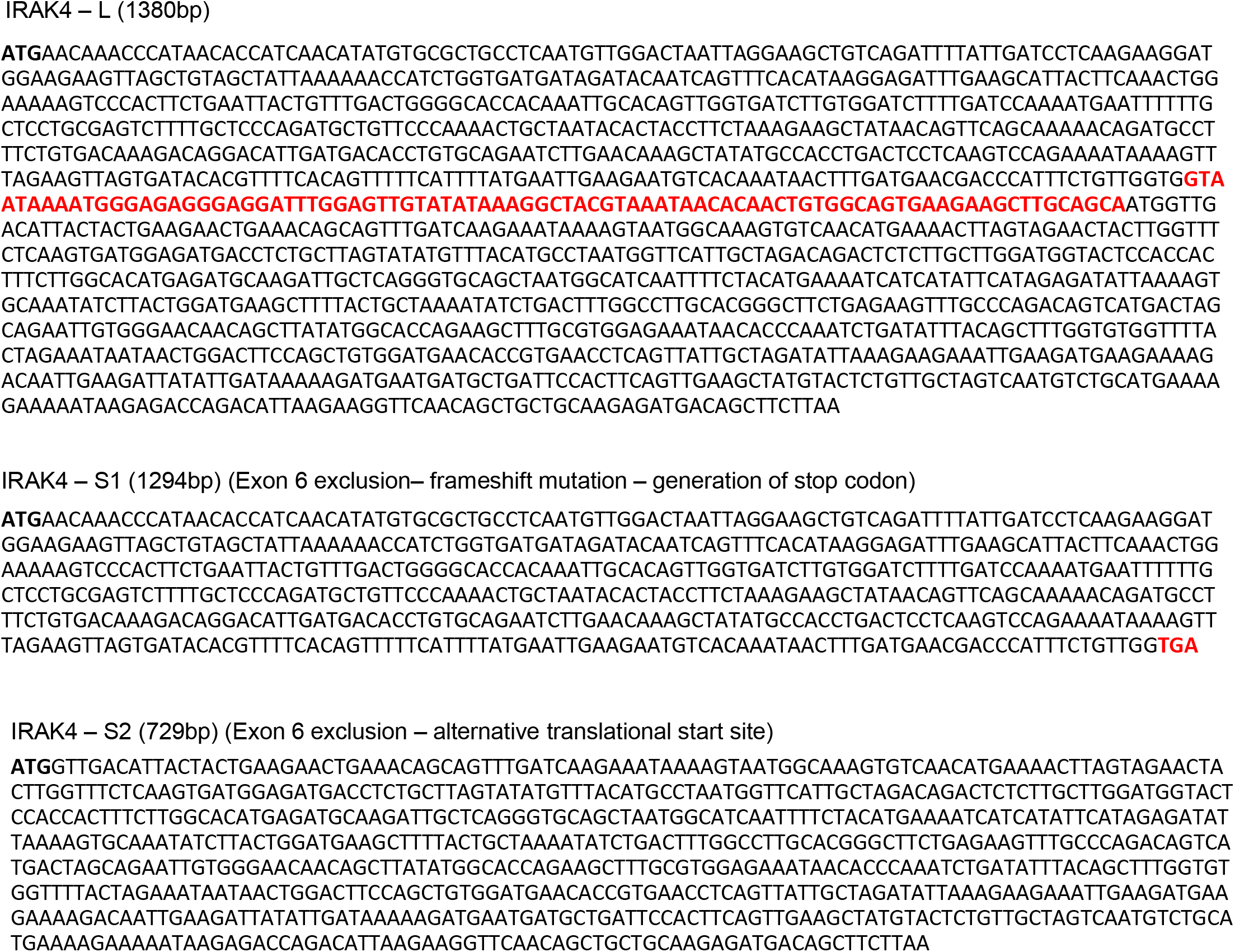
Amino acid sequences of IRAK4-Long and the shorter IRAK4-Sequence 1 and IRAK4-Sequence 2 isoforms: The amino acids shown in red color (exon 6) are retained in IRAK4 Long isoform.

**Supplementary Figure 2:**
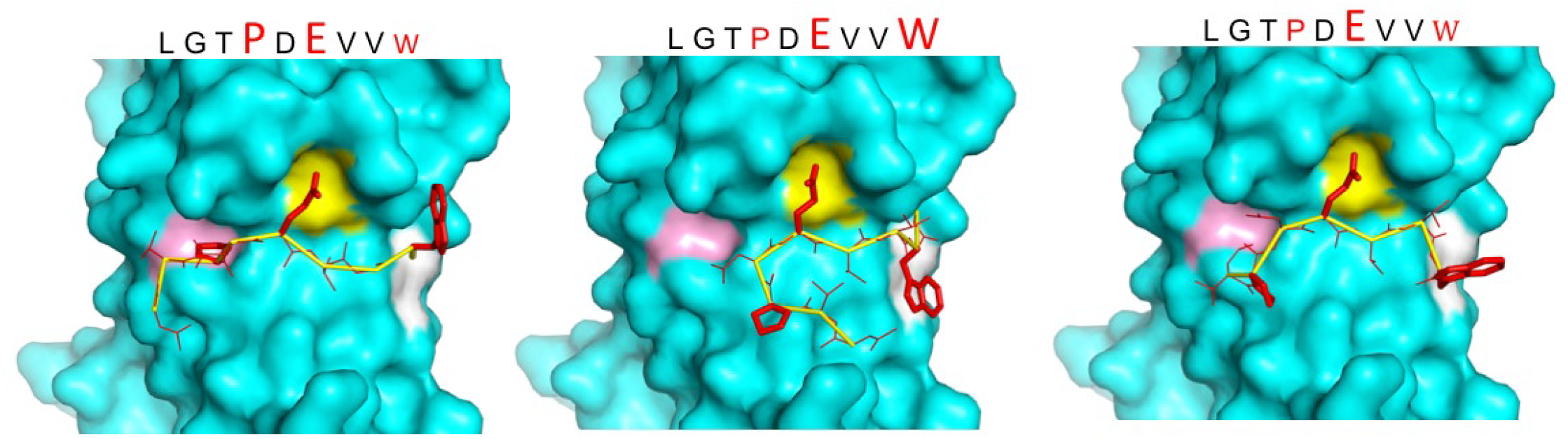
Modelling interaction between TRAF6 and CDK2: MMGBSA docking (see Methods) of CDK2 peptide to substrate binding domain of TRAF6 (PDB: 1LB5). Various potential clusters are shown. TRAF6 and CDK2 peptide are shown in ribbon and surface representation, respectively. High-lettering of conserved residues P, E, W within the shown peptides depicts successful docking in corresponding sub-pockets.

**Supplementary figure 3:**
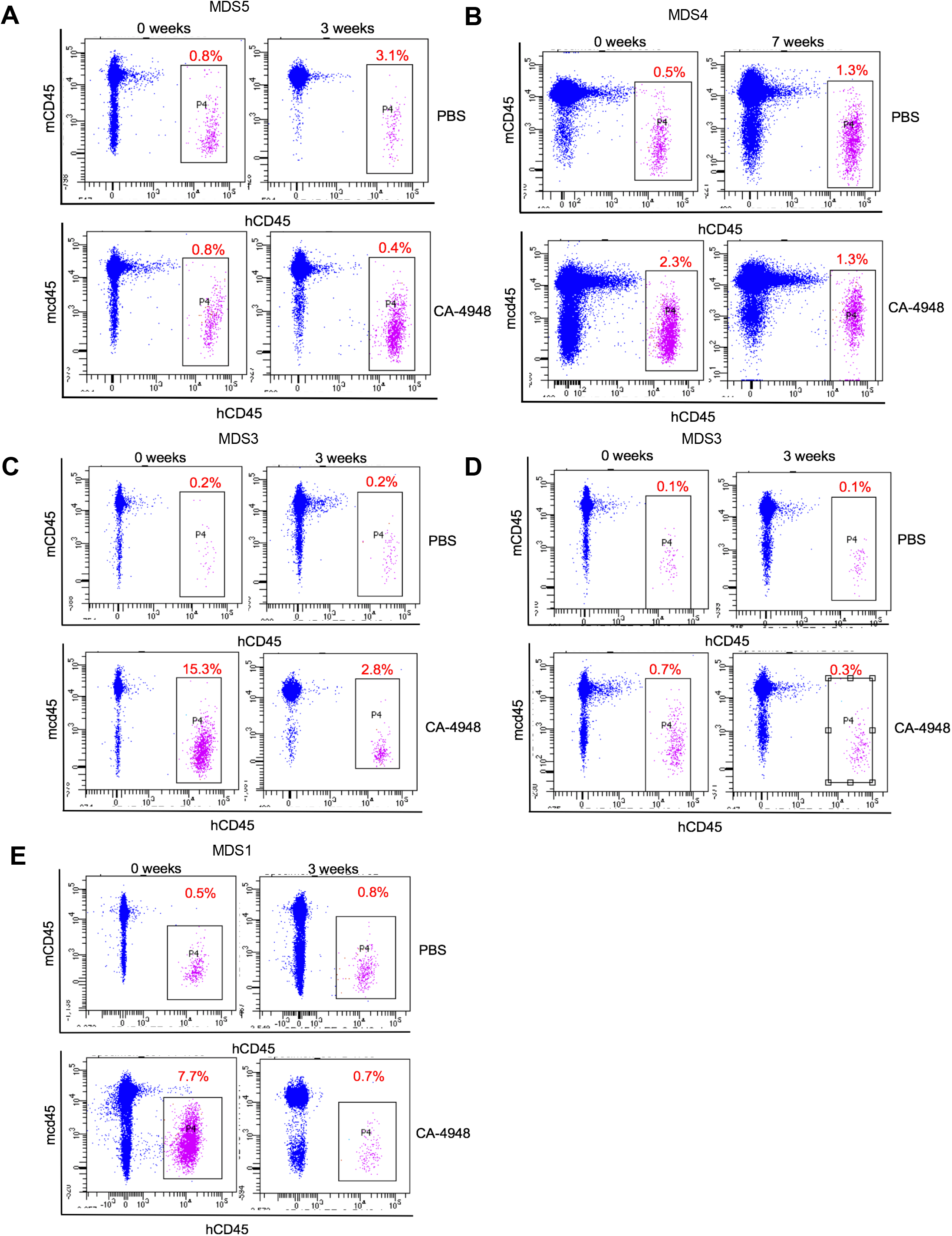
IRAK4 inhibition leads to reduction in MDS clones in vivo: NSG mice were xenografted with MDS cells with SF3B1 mutation. After engraftment, mice were treated with either vehicle or IRAK4 inhibitor (CA-4849, 12.5 mg/kg) and bone marrow aspirates were used to evaluate human cells engraftment by flow cytometry.

**Supplementary figure 4:**
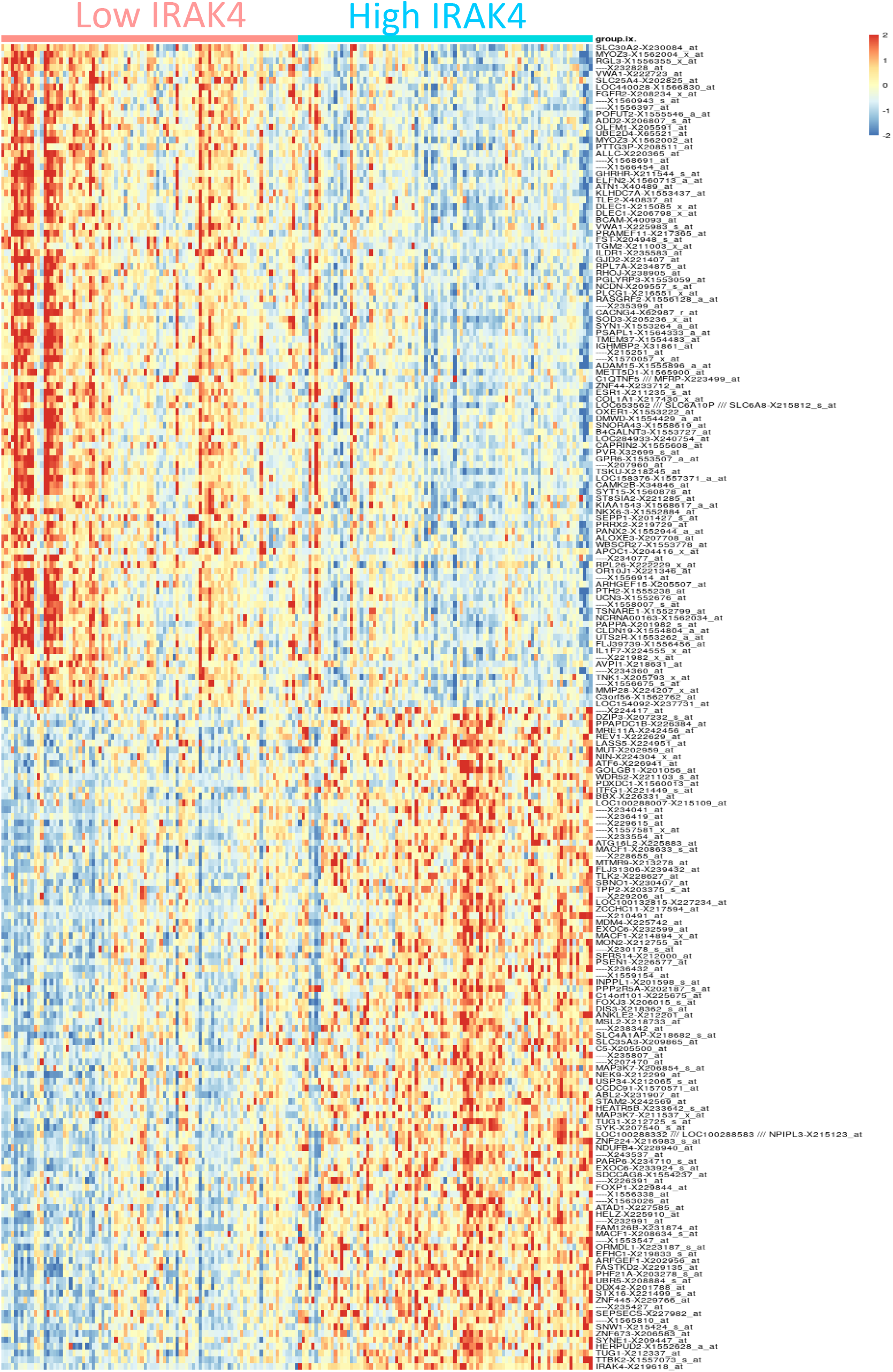
Gene expression signature associated with high IRAK4 expression. Top 100 differentially expressed genes between cohorts with high and low IRAK4 (based on median expression) are shown.

**Supplementary figure 5:**
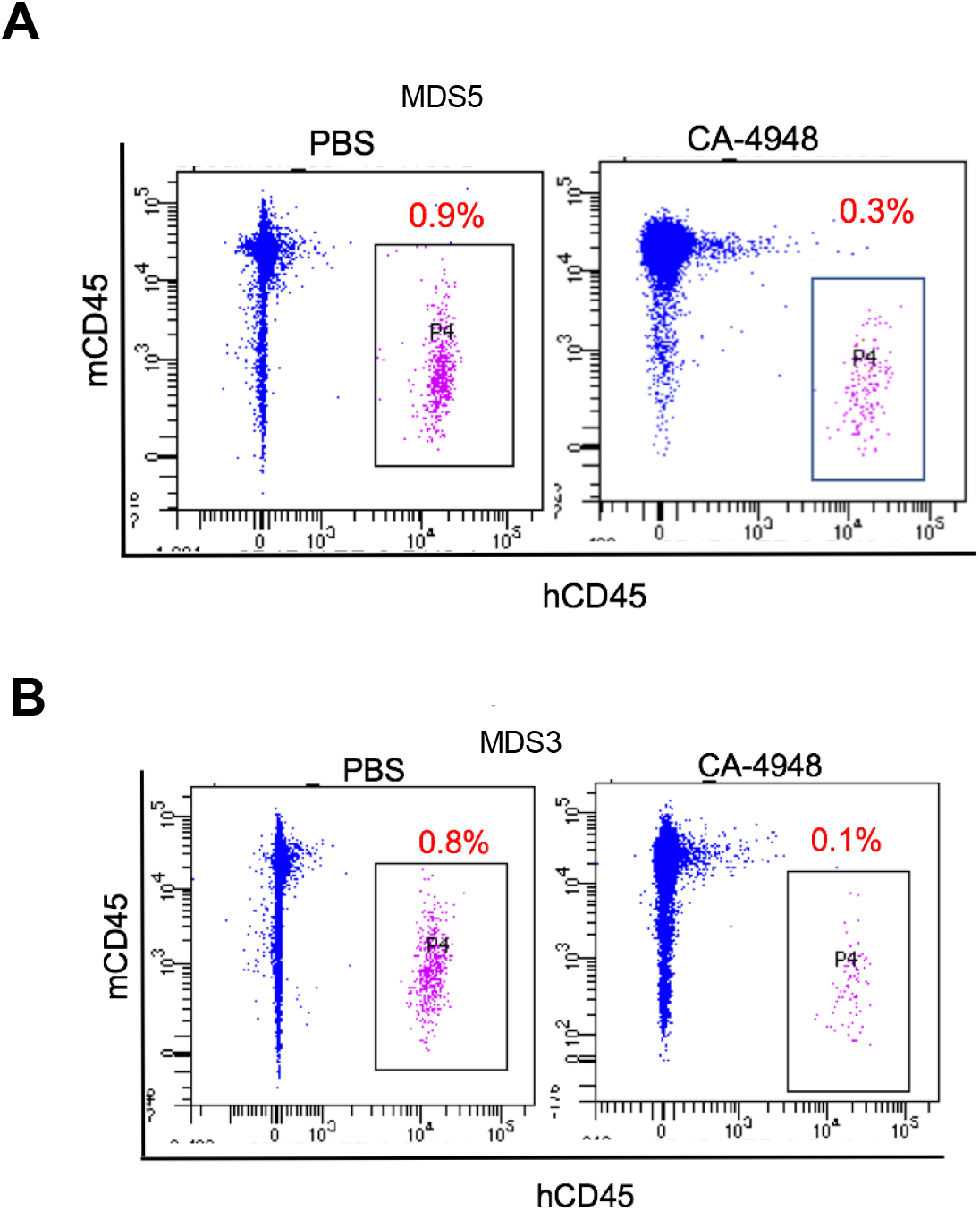
IRAK4 inhibition leads to reduction in MDS clones in secondary transplant recipient mice: MDS bone marrow cells were isolated and purified from NSG mice either treated with vehicle or CA-4948 in and then xenografted in secondary recipient NSG mice. Human cell engraftment was evaluated after 4 weeks by flow cytometry on bone marrow aspirates

**Supplementary figure 6:**
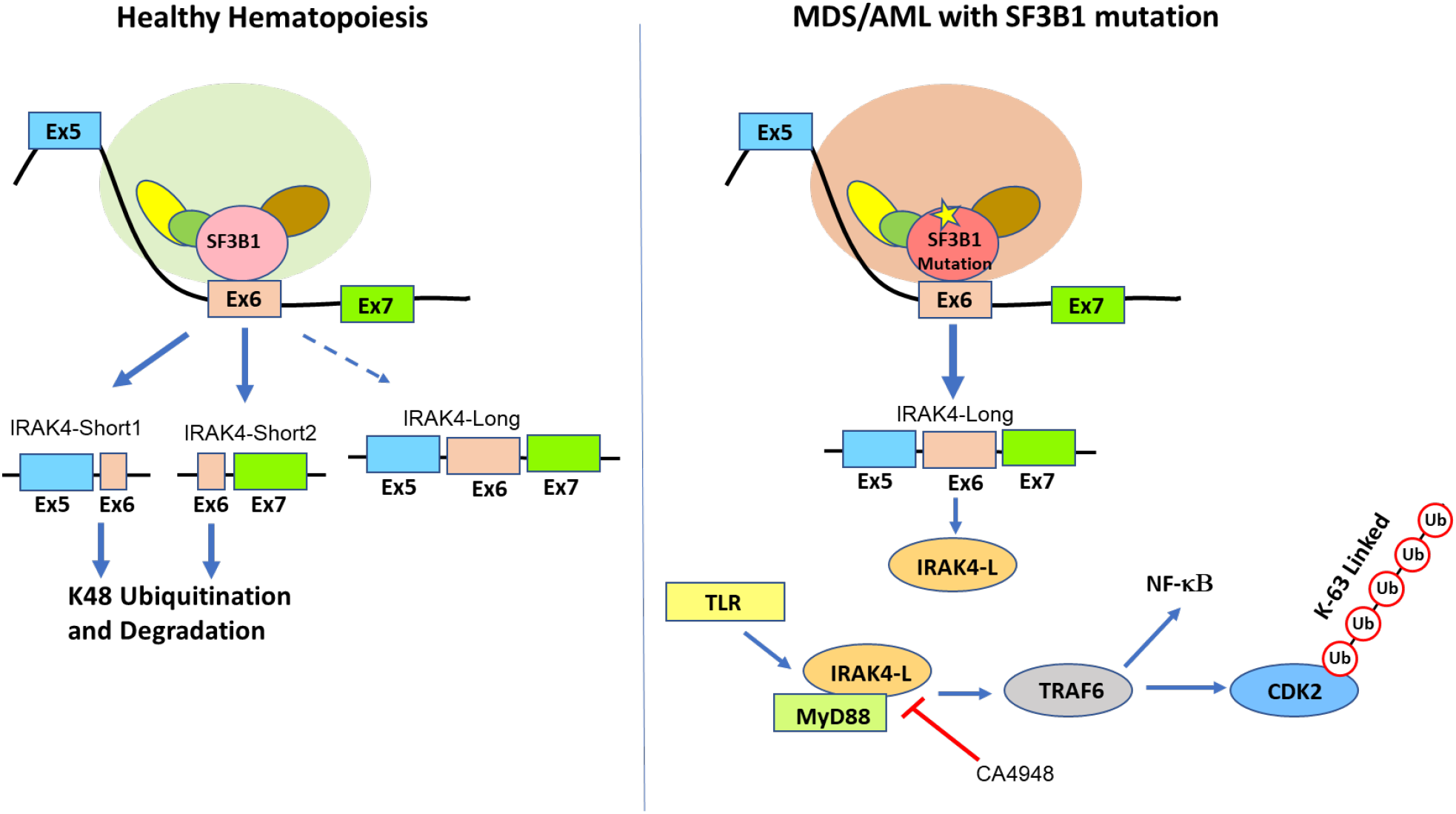
Proposed schematic of MDS pathogenesis due to missplicing of IRAK4 because of SF3B1 mutations

**Supplementary Figure 7:**
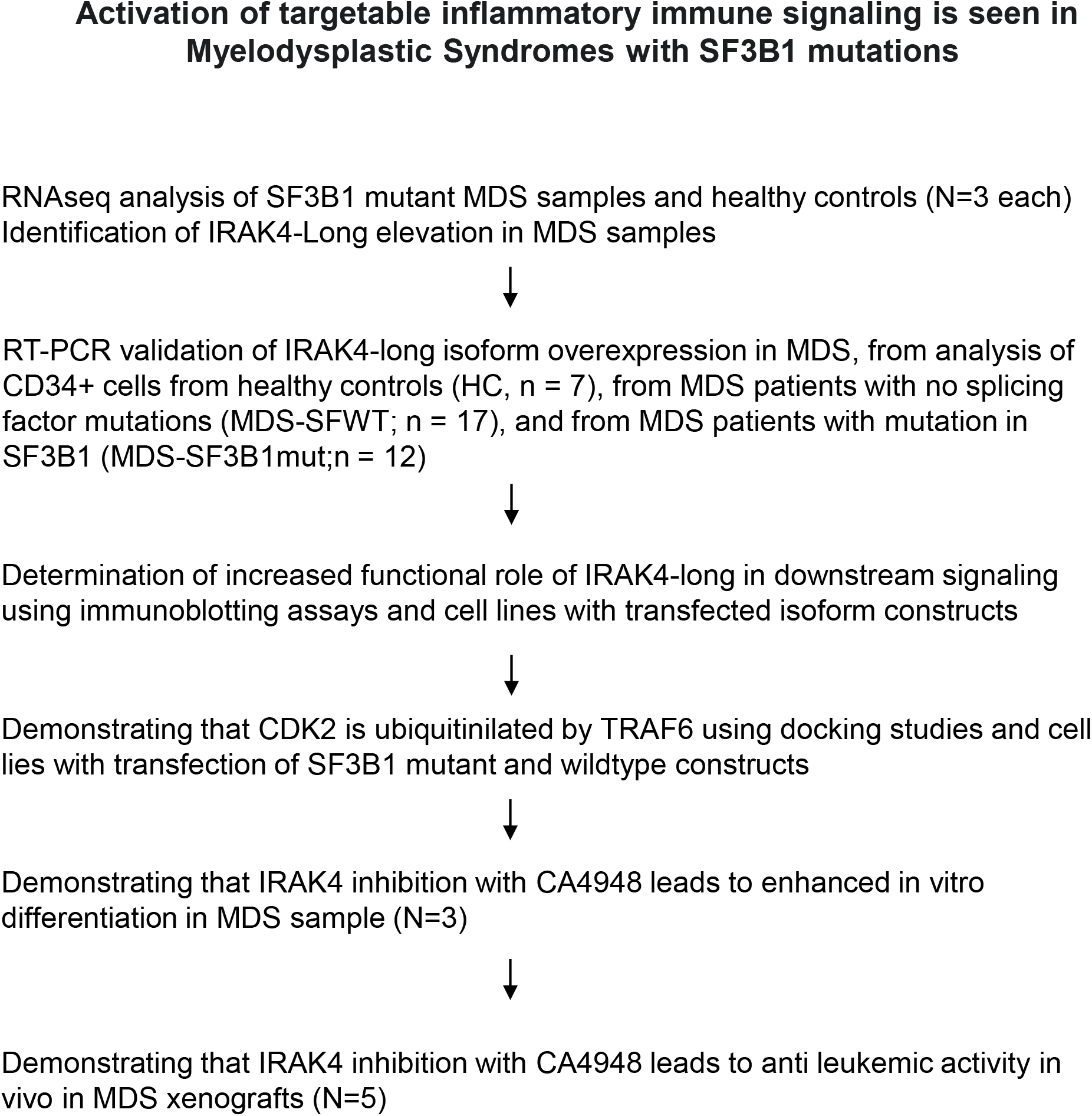
STROBE flowchart showing study schema

**Supplementary Table 1:** Patient samples.

| Diagnosis | Subtype | Mutations | Hgb<br>(GM/DL) | WBC<br>(X10<sup>3</sup>/ML) | Plts<br>(X10<sup>3</sup>/ML) | Cytogenetics |
| --- | --- | --- | --- | --- | --- | --- |
| MDS1 | RA | SF3B1, TET2 | 8 | 6 | 110 | NML |
| MDS2 | RAEB | SF3B1, RUNX1,<br>BCOR, NRAS,<br>DNMT3A | 6 | 2 | 60 | +13 |
| MDS3 | RCMD | SF3B1, IDH1, JAK2,<br>KRAS, TET2 | 7 | 3 | 50 | NML |
| MDS4 | RCMD | SF3B1, TET2 | 8 | 5 | 120 | del(5q) |
| MDS5 | RCMD | SF3B1 | 7 | 1 | 30 | NML |
| MDS6 | RCMD | SF3B1, TET2,<br>GNAS | 7 | 3 | 90 | NML |

